# Deciphering cis-regulatory elements using REgulamentary

**DOI:** 10.1101/2024.05.24.595662

**Authors:** Simone G. Riva, Emily Georgiades, Jennifer C. Herrmann, E. Ravza Gür, Edward Sanders, Martin Sergeant, Matthew Baxter, Jim R. Hughes

**Affiliations:** MRC Molecular Haematology Unit, MRC Weatherall Institute of Molecular Medicine, University of Oxford, Oxford, UK; MRC WIMM Centre for Computational Biology, MRC Weatherall Institute of Molecular Medicine, University of Oxford, Oxford, UK

**Keywords:** Bioinformatics, gene regulation, cis-regulatory elements

## Abstract

With the boom in Genome-Wide Association Studies (GWAS), it has become apparent that many disease-associated genetic variants lie in the non-coding regions of the genome. In order to prioritise these variants and disentangle their functional significance, it is important to be able to accurately classify cis-regulatory elements within these non-coding regions of the genome. Historically, the classification of cis-regulatory elements relied purely on the presence of characteristic histone marks, with recent advancements in their classification using more sophisticated Hidden Markov Model (HMM)-based approaches. The limitation of the HMM-based approaches is that the output of these models is an arbitrary chromatin state, which then requires the user to manually assign these states to a particular class of cis-regulatory elements. Here we present a new tool, REgulamentary, which enables *de novo* genome-wide annotation of cis-regulatory elements in a cell-type specific manner. We benchmarked REgulamentary against GenoSTAN, the most popular existing published chromatin annotation and regulatory element identification tool, to demonstrate the advancements REgulamentary can provide in assigning chromatin states. Finally, as an example of REgulamentary’s utility in solving complex disease trait *loci*, we applied REgulamentary to published GWAS data to demonstrate how this tool can be used to prioritise likely causal variants.

## 1. Introduction

The non-coding genome is known to be populated with thousands of regulatory elements which act to control spatiotemporal gene expression. Two of the most important classes of regulatory elements are enhancers and promoters which have specific but overlapping roles. Promoters act to enable transcription initiation, they are typically short sequences, located close to the transcription start site (100bp-1kb). Active promoters, that is those associated with actively transcribed genes, reside in regions of open chromatin, and display the characteristic tri-methylation of Histone H3 at Lysine 4 (H3K4me3). Promoter elements contain general transcription factor binding motifs and thus act as a platform on which a compendium of transcription factors can bind and associate with general transcription machinery in order to initiate transcription. Enhancer elements on the other hand, are located more distally from the target gene, at distances of up to 1Mb in some cases. Enhancer sequences are known to encode the cell type-specific transcription factor binding sites. When active, enhancers exhibit the characteristic histone signatures: mono-methylation of Histone H3 at Lysine 4 (H3K4me1) and acetylation of the lysine residue at N-terminal position 27 of the histone H3 (H3K27ac), and recruit tissue-specific transcription factors in order to regulate cell type-specific gene expression [1, 2].

The classical definitions of enhancers and promoters stated above are somewhat oversimplified. It is well known that there is a high degree of overlap between the histone modifications displayed by active enhancers and promoters. It is not possible to predict which target genes an enhancer will interact with by genomic position alone, enhancers often skip the most proximal genes and may interact with more than one gene within a given locus, thus metrics such as distance to the nearest transcription start site (TSS) can often be misleading. For many purposes it is of interest to determine the activity of cis-regulatory elements, for example, to know whether they are active in a given cell type. To directly measure the activity of a cis-regulatory element is challenging, therefore it is common practice to infer the activity of an element based on chromatin accessibility, associated histone marks, and transcription factor binding. However, given the degree of overlap in these classifiers across enhancers and promoters, this is not a simple task, and depending on the chosen method this can give varying results [1, 2].

Active regulatory elements are located in regions of open chromatin and marked by specific features, which can be used to putatively identify them in the genome. H3K4me3 and H3K27ac are associated with active promoters, whereas H3K4me1 and H3K27ac are found at active enhancers [1, 2, 3, 4]. Boundary elements are an additional class of cis-regulatory elements, which in contrast, are predominantly characterised by the binding of the CCCTC-binding factor (CTCF) [3, 5]. To classify regulatory elements within any given eukaryotic cell type, REgulamentary therefore requires: (1) the positions of regions of open chromatin based on either DNase-seq [6], ATAC-seq [7] or single-cell ATAC-seq [8], (2) genome-wide data for the histone modifications H3K27ac, H3K4me1 and H3K4me3 based on ChIP-seq, and (3) genome-wide data for CTCF based on ChIP-seq [9, 10]. Further details on the input files required for REgulamentary can be found in the Methods section.

There have been a number of bioinformatic tools built for the task of cis-regulatory element classification, however, there is a lack of a substantial ground-truth dataset in which both enhancer and promoter elements have been experimentally validated. In addition, the identification of regulatory elements is data-driven, high-quality and cell-type-specific input data is required in order to achieve an accurate classification. Unsupervised learning methods might be suited to this task, however these methods are not able to assign regulatory elements, they instead group them by their similarities. For this reason, we opted for a more systematic approach. In this work, we propose REgulamentary, a rule-based model built to identify the regulatory elements. REgulamentary takes into account the characteristics of each element, as they would be manually assigned by an expert biologist, in an automated and genome-wide manner. We demonstrate that our proposed approach is able to outperform GenoSTAN, the current most widely used cis-regulatory element classification tool available.

## 2. Method

REgulamentary takes the following input data types: (1) chromatin accessibility data (this can be either ATAC, scATAC, or DNase), (2) three histone mark ChIP-seq data sets (H3K4me1, H3K4me3, and H3K27ac), and (3) ChIP-seq for the CTCF boundary element. For data types (2) and (3), aligned sequences (bam format), coverage (bigwig format), and peak files (bed format) are required, whereas for chromatin accessibility only the peak bed file is necessary. Table 1 shows the data that has to be provided to REgulamentary.

**Table 1.**
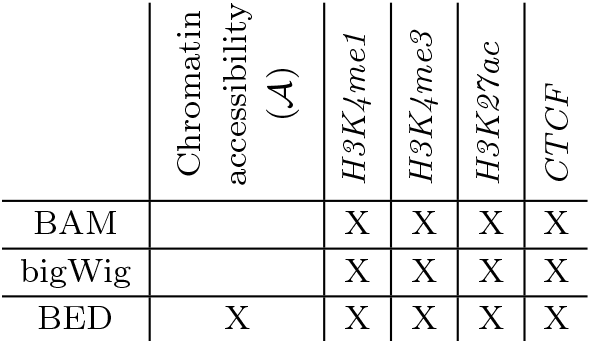
Table shows the data formats required as input to REgulamentary. Boxes marked with X are required data types. BAM, bigWig, and bed files for each target must originate from each other, e.g., the CTCF BED must be a bed file containing peaks called from the provided CTCF bigwig file, where peaks are naturally variable in size.

|  | Chromatin accessibility (A) | H3K4me1 | H3K4me3 | H3K27ac | CTCF |
| --- | --- | --- | --- | --- | --- |
| BAM |  | X | X | X | X |
| bigWig |  | X | X | X | X |
| BED | X | X | X | X | X |

### 2.1. Step 1 - Data pre-processing

For pre-processing and meta-plot visualisation, REgulamentary concatenates, sorts by coverage in ascending order, and finds the unique peaks between 𝒜 and *CTCF* peak files, to create the list of union peaks (*<u>r</u>*). From *<u>r</u>*, peaks intersecting with blacklist regions are then removed. Using the multi-coverage function within bedtools [11], a **C**_*r*_ coverage matrix is calculated for each *r* ∈ *<u>r</u>* for *H3K4me1* and *H3K4me3* histone mark ChIP-seq data. Within **C**_*r*_, each *r* ∈ *<u>r</u>* is then ranked in descending order based on the difference between the coverage values of *H3K4me3* and *H3K4me1*.

### 2.2. Step 2 - Metric generation

Let *auc* (Area Under the Curve — read count for provided regions) [12], be a function defined as follows: *auc* : *<u>r</u> → ℕ* _*≥*0_, where *auc*(*r*) := read coverage of *r*. After data pre-processing, REgulamentary intersects *<u>r</u>*, the regions of interest, with the three histone mark ChIP-seq and the CTCF boundary element and uses *auc* to calculate the normalised — per genome-wide sequencing depth — read counts, creating a *R* × *T* matrix 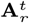 (eq. 1), where *R* = |*<u>r</u>*| and *T* = 4, which represents the four tracks: *<u>t</u>* = [*H3K4me1, H3K4me3, H3K27ac, CTCF* ]. 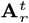 is defined as:

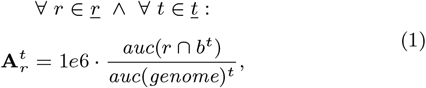

where *b*^*t*^ indicates the regions of interest for *t* ∈ *<u>t</u>*.

### 2.3. Step 3 - Rank

Let *rank* be a function defined as follows: *rank* : 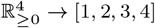. For each *r* ∈ *<u>r</u>*, 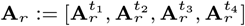 is ranked in descending order, accordingly to **A**_*r*_’s values, creating a *R* × *T* matrix 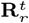 where (eq. 2)^*^:

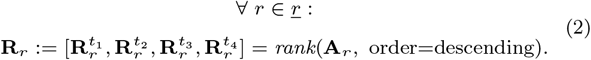

### 2.4. Step 4 - Annotation

Finally, REgulamentary systematically assigns to each *r* ∈ *<u>r</u>* a regulatory element (RE) label (namely Enhancer, Promoter, CTCF, Enhancer/CTCF, and Promoter/CTCF) in two phases. The first phase assigns the main RE: Enhancer, Promoter, and CTCF, by applying the following rule (3)^*^:

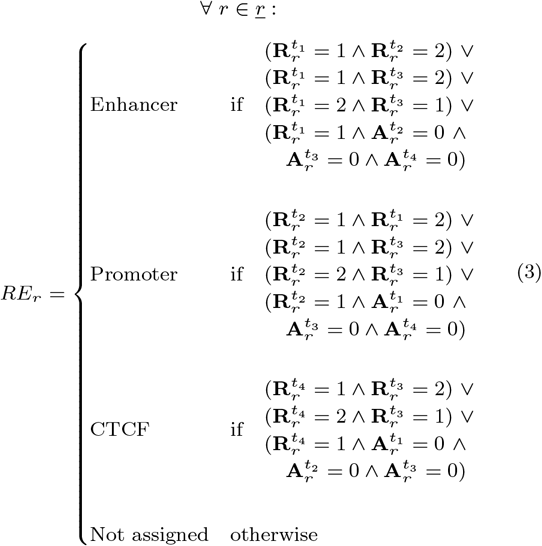

whilst the second phase tries to discriminate Enhancer or Promoter regions which are co-accessible with CTCF sites (Enhancer/CTCF and Promoter/CTCF), by applying (4)^*^:

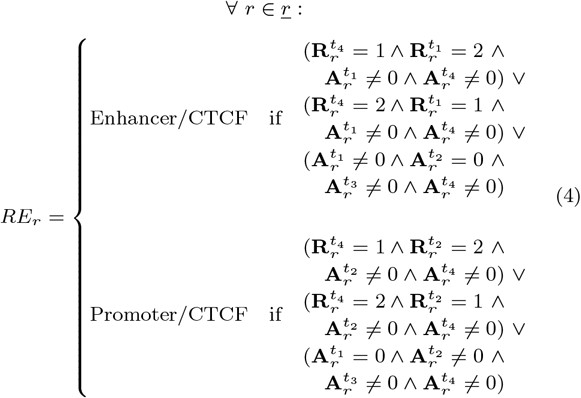

## 3. Results

From ENCODE we downloaded publicly available FASTQ files for: DNase-seq (n=14), ChIP-seq of the histone modifications H3K27ac (n=2), H3K4me1 (n=3) and H3K4me3 (n=7) and ChIP-seq of CTCF (n=7) in human umbilical vein endothelial cells (HUVECs) [13] (see Supplementary Section A for ENCODE accession sample ids). The FASTQ files were first processed using CATCH-UP [14], a portable automated pipeline for analysing bulk ATAC-seq and ChIP-seq data. In addition to BAM files, CATCH-UP outputs bigWig files for visualisation and BED files containing the peak calls for each sample, which can be readily used as input for REgulamentary.

### 3.1. REgulamentary result

As stated in Section 2.1, all regions of interest (total of 33792 sites) intersect chromatin accessibility regions from DNase-seq, and have variable CTCF coverage.

Figure 2a shows the 33792 peak regions of interest, derived from the processed HUVEC DNase-seq data. These regions of interest were then assigned to specific classes of regulatory elements using REgulamentary, and are grouped together in Figure 2b. Out of the 33792 regions of interest, 33669 (*>* 99.996%) were assigned as regulatory elements, demonstrating the thoroughness with which REgulamentary assigns a regulatory element to each region of interest. The most commonly identified regulatory elements were CTCF site (33.3%), followed by Enhancers (28.3%) and Promoters (24.7%), as shown in Figure 3. A small fraction of the regions showed characteristics of either a promoter or an enhancer that overlapped with CTCF binding (10.5% and 3.3%, respectively).

**Fig. 1:**
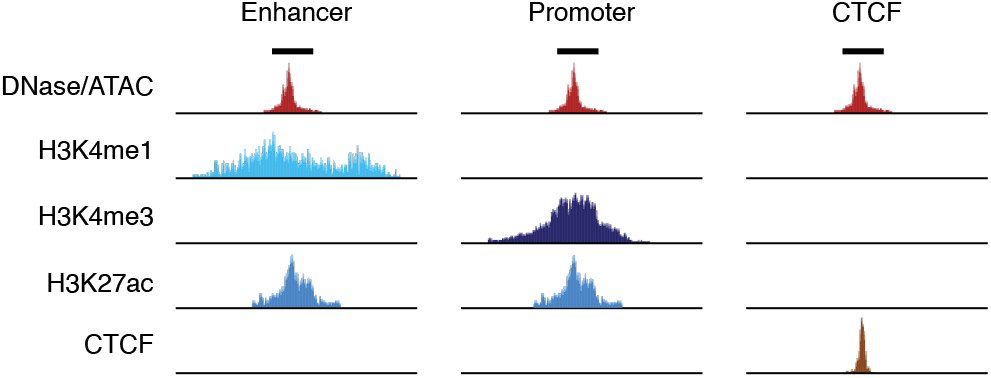
Visualisation of how regulatory elements are identified by using chromatin accessibility data (such as DNase, bulk-ATAC, or single-cell ATAC-seq data), H3K4me1, H3K4me3, H3K27ac histone modification marks, and ChIP CTCF-seq data.

**Fig. 2:**
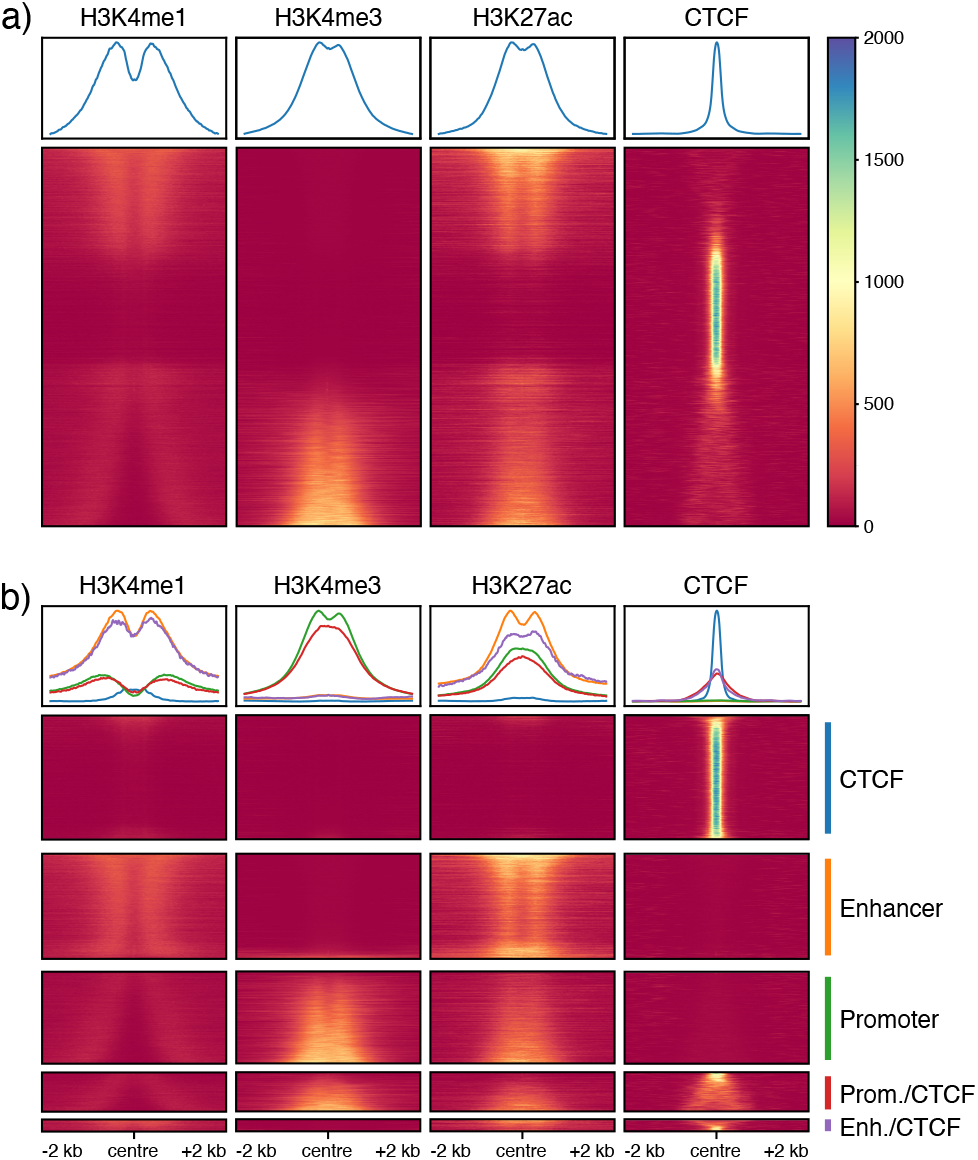
In a) is shown the meta-plot of the coverage *±*2kb from the centre of the sorted regions of interest in HUVECs (as explained in 2.1), whilst b) shows the same meta-plot grouped instead by regulatory element assigned by REgulamentary.

**Fig. 3:**
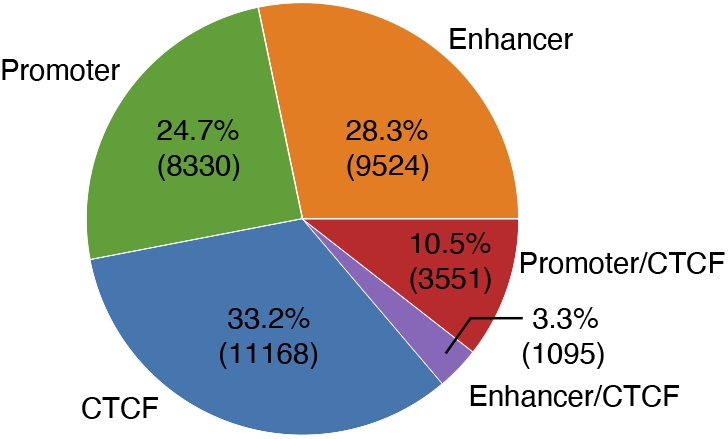
The piechart displays the distribution of regulatory elements in HUVEC regions of interest.

### 3.2. Benchmarking against GenoSTAN

In order to benchmark REgulamentary, we compared the output to the current gold standard tool for genome annotation, Genomic STate ANnotation (GenoSTAN) [15]. GenoSTAN is a genome segmentation method that is available as part of R/Bioconductor package STAN [16]. It relies on learning chromatin states from a sequence of samples by performing Hidden Markov Models (HMM) with flexible multivariate count distributions. For comparison purposes, we initialise and fit (bidirectionally) GenoSTAN with 6 states and use Gaussian as the emission distribution of the model. For each region of interest, we calculate the most likely state path using Viterbi. We choose 6 states to have a fair comparison with the 6 possible identities REgulamentary can assess (‘Not assigned’ included). After the assignment of the chromatin states, we used a heatmap plot (Figure 4), showing the normalised read counts per state for H3K4me1, H3K4me4, H3K27ac, and CTCF to assess the regulatory elements, which have been manually assigned based on visual assessment (Table 2) to each GenoSTAN state, according to the normalised read counts and according to the naming automatically given by REgulamentary.

**Table 2.** Manual assessment of GenoSTAN states, based on (log) scaled read counts (Figure 4), according to the REgulamentary naming assignment.

| State | Annotation |
| --- | --- |
| 1 | Not assigned |
| 2 | Promoter |
| 3 | CTCF |
| 4 | Enhancer |
| 5 | Promoter/CTCF |
| 6 | Promoter |

**Fig. 4:**
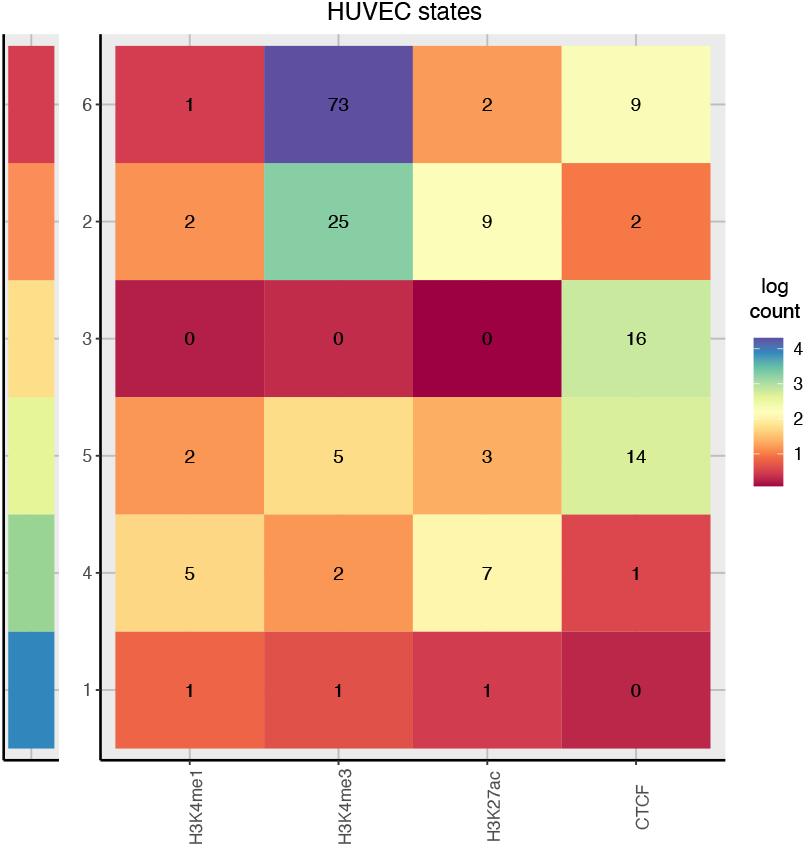
The heatmap shows the distribution of log normalised count for each GenoSTAN state across H3K4me1, H3K4me4, H3K27ac, and CTCF.

As a first visual result, we re-plot the meta-plot for comparison purposes, this time grouping regions of interest based on GenoSTAN state annotation in Figure 5, where it is possible to see that some regions were miss-classified by using the GenoSTAN approach. For example, the Promoter/CTCF group contains a significant subset of regions with a high coverage of H3K4me1, a mark characteristic of enhancer classes, suggestive of a miss-classification.

**Fig. 5:**
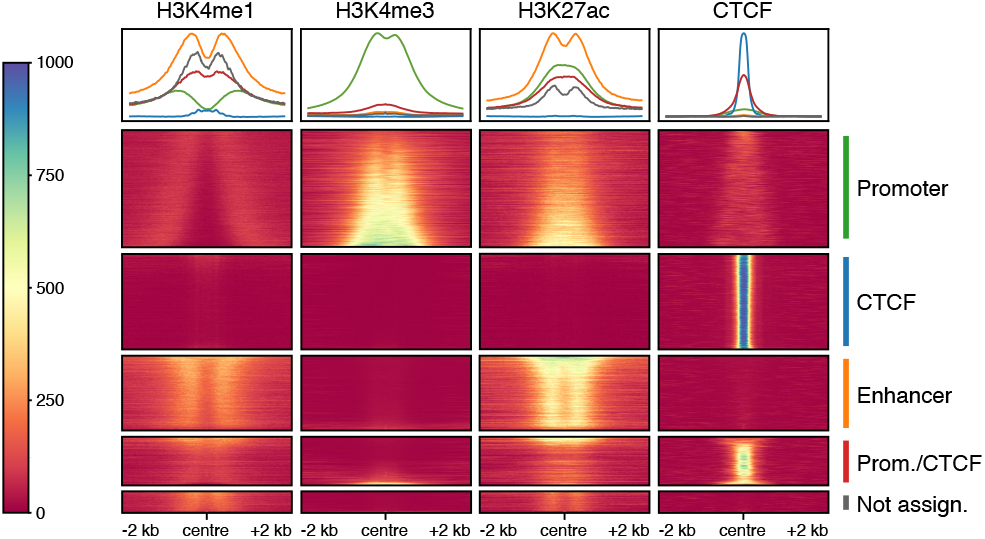
The meta-plot displays the coverage *±*2kb from the centre of the sorted (as explained in 2.1) regions of interest in HUVECs grouped by the manual assignment (Table 2) of the regulatory elements to the GenoSTAN states.

In order to compare REgulamentary against GenoSTAN, it is important to understand how closely the classifications of each of the 33792 regions of interest agree (Figure 6). First, we show that for 2715 regions the output of the two tools closely align: REgulamentary identifies as Promoter/CTCF and GenoSTAN as Promoter. However, approximately a third of the total regions of interest (9895 out of 33792) were classified differently by REgulamentary and GenoSTAN. Of interest are the 300 regions that REgulamentary identifies as Promoter and GenoSTAN identifies as Enhancer, representing a high degree of mismatch between the two methods. This mismatch is also observed in the 538 Enhancer-Promoter (REgulamentary-GenoSTAN, respectively), and the 132 CTCF-Enhancer (REgulamentary-GenoSTAN, respectively). In these cases, we randomly selected an example of each of these classes to manually investigate these discrepancies. For example, the promoter of TUBA4A, a gene which encodes a key component of cytoskeletal microtubules, is correctly assigned by REgulamentary, but incorrectly assigned as an enhancer by GenoSTAN (Figure 7a). This promoter exhibits both H3K4me1 and H3K4me3 signals, which may explain the discrepancy. However, REgulamentary is able to correctly identify this feature due to the correct ranking of the relative signal strength of the two chromatin marks. Similarly, an enhancer located downstream of the vascular endothelial surface protein PCDH1 gene which exhibits high levels of H3K4me1 and low levels of H3K4me3 is correctly identified by REgulamentary, but misidentified as a promoter by GenoSTAN (Figure 7b). Finally, a CTCF site located in an intron of the LSG1 gene, which also exhibits some diffuse nearby H3K4me1 signal, again, is correctly identified by REgulamentary but misassigned as an enhancer by GenoSTAN (Figure 7c).

**Fig. 6:**
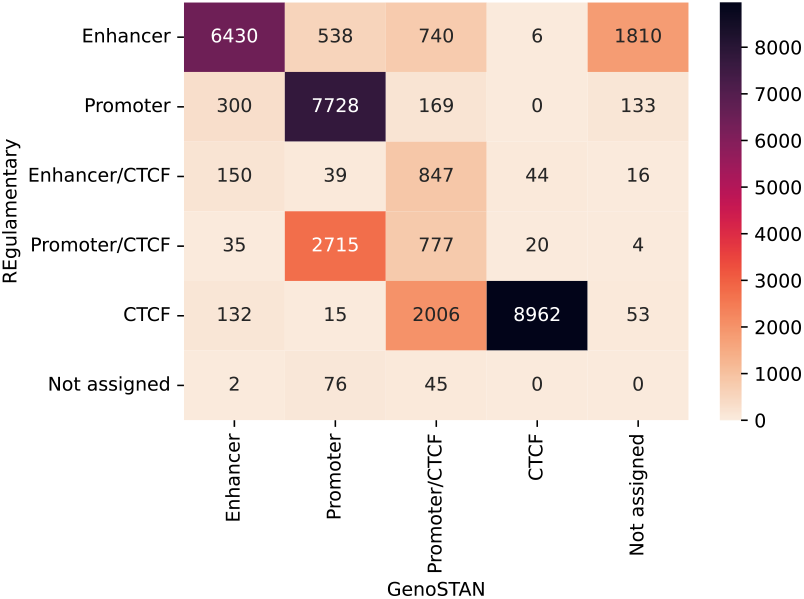
The confusion matrix displays the comparison in absolute numbers of the regulatory elements between REgulamentary (y-axis) and GenoSTAN (x-axis) outputs, respectively.

**Fig. 7:**
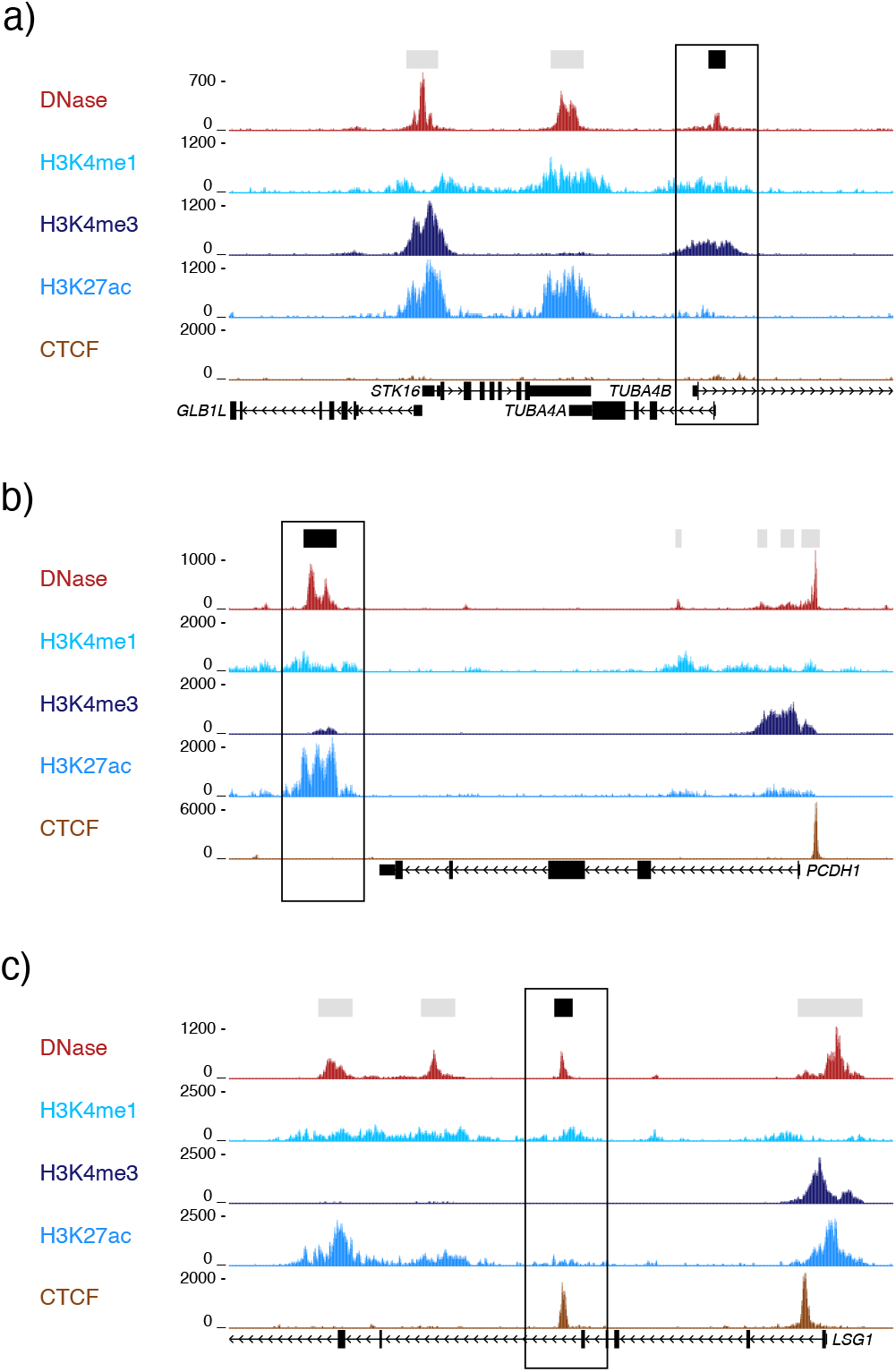
Highlighted, in a) is shown a correctly assigned Promoter region by REgulamentary but misassigned as Enhancer by GenoSTAN. Likewise for b), a correct Enhancer region for REgulamentary and misassigned by GenoSTAN as Promoter. Lastly, in c), a properly annotated CTCF region by REgulamentary and misassigned Enhancer by GenoSTAN.

### 3.3. Intersection with stroke GWAS

Prioritisation of the common non-coding genetic variants found in GWAS requires the accurate identification of cell-type-specific regulatory elements. To demonstrate the utility of REgulamentary, we selected and downloaded stroke-related lead SNPs from the GWAS Catalog [17], version 1.0.2, downloaded on 22/08/2023 (see table Supplementary Section B for a comprehensive list), and imputed proxy SNPs, *R*^2^ filtered (*R*^2^ ≥ 0.8), by using TopLD [18] on the European population. Stroke is a leading cause of morbidity and mortality in the world, with a significant genetic component, and represents a large unmet clinical need. The underlying pathological mechanisms are thought to involve endothelial cells, therefore we predict that a proportion of the genetic associations will be active in HUVECs [19, 20]. In total, 17571 stroke-associated imputed SNPs were intersected with the REgulamentary defined HUVEC regulatory elements. 94 SNPs were identified in Enhancers, along with 85 in promoters, and 65 in CTCF elements. A further 19 and 30 were identified in Enhancer/CTCF and Promoter/CTCF elements (Figure 8). The assignment of the elements containing these SNPs, differed significantly when the same analysis was run using GenoSTAN annotation of HUVECs, demonstrating the importance of an accurate annotation of cell-type-specific regulatory elements. Accurate prioritisation of disease-associated genetic variants may expedite the understanding of complex diseases and the search for novel therapies.

**Fig. 8:**
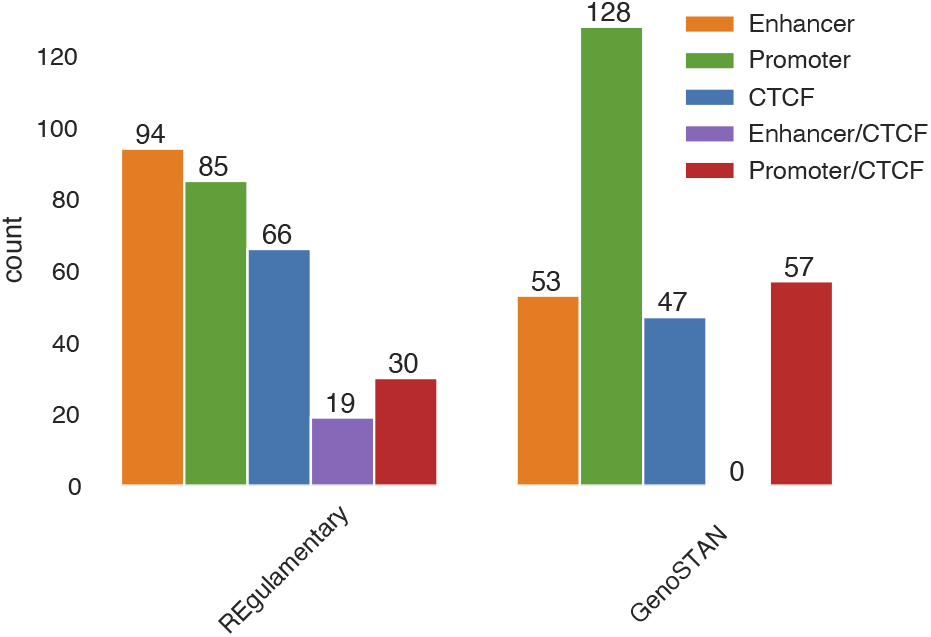
The barplot shows the number of stroke GWAS genetics intersection with REgulamentary and GenoSTAN annotated regulatory elements, respectively.

## 4. Conclusion

In this work, we presented REgulamentary, a standalone, rule-based bioinformatic tool for the thorough annotation of cis-regulatory elements for chromatin-accessible or CTCF-binding regions of interest. We showed that the proposed tool is able to classify regions of interest accurately (Figure 2b) and outperforms the current gold-standard tool — GenoSTAN — (Figure 6), showing a direct comparison between the tools (Figure 7).

Whilst we have shown our tool is able to surpass the current gold-standard approach, we would like to highlight four key areas which will form the focus of future developments of this tool: (1) first, it is widely known that directly measuring the activity of a cis-regulatory element is challenging, therefore it is common practice to infer the activity of an element based on chromatin accessibility, associated histone marks, and transcription factor binding. However, given the degree of overlap in these classifiers across enhancers and promoters, this is a complex task, and depending on the chosen method this can give varying results as we have demonstrated. It is known that specific compendiums of transcription factors bind at either promoter or enhancer elements, in a specific spatio-temporal manner. We therefore propose providing a list of ChIP-seq transcription factors peaks and intersecting these with the output annotations of REgulamentary to enhance the accuracy and biological significance of the output, by resolving the classifications for specific cell types or differentiation stages. In doing so the output of REgulamentary would have the ability to classify the regulatory elements by the activity of the elements, given transcription factor binding can be used as a proxy for this. (2) Secondly, we would like to increase the speed and efficiency of the pipeline. The current version handles read counts, normalization, and *auc* computation in parallel using Python’s multiprocess feature, which despite being streamlined still requires significant run-time (∼ 5h) based on the size of the input data). To address this issue, we aim to implement Graphics Processing Units (GPUs) to increase the efficiency of data processing [21], and thereby speed up (∼ 10X) the assessment of cis-regulatory elements. (3) Thirdly, we aim to use REgulamentary to create a robust and reliable roadmap of regulatory elements, covering as many cell types as possible. This will create a first-in-class dataset which can be used to train, validate, and test Deep Learning (DL) Models [22], such as Convolutional Neural Networks [23], Recurrent Neural Networks [24], or Transformers [25]. This dataset will provide sufficient positive and validated examples of regulatory element regions per cell type in a suitable format for DL approaches, which will eliminate the need for histone marks and CTCF ChIP-seq input data, relying instead only on chromatin accessibility data to identify the different cis-regulatory elements accurately. (4) Lastly, by creating a WebApp for REgulamentary, it will become more user-friendly and accessible to a broader range of scientists.

## Supporting information

Supplementary A - ENCODE Accession IDs

Supplementary B - Lead SNPs

## Code availability

REgulamentary is a Python-based Snakemake [26] pipeline built to identify cis-regulatory elements from sequenced data (GitHub: github.com/Genome-Function-Initiative-Oxford/REgulamentary).

## Acknowledgments

S.G.R. is supported by the MRC grant (MC UU 00029/3). E.G. is supported by the Wellcome Genomic Medicine and Statistics PhD Programme (108861/Z/15/Z) and the MRC grant (MC UU 00029/3). J.C.H. is supported by the Medical Research Council (MC UU 00016/14 and MR/N00969X/1) and the Wellcome Trust grants (106130/Z/14/Z and 108861/B/15/Z). E.R.G. is supported by the Ministry of National Education Selection and Placement of Candidates Sent Abroad for Postgraduate Education (YLSY) scholarship, Republic of Türkiye Ministry of National Education and the Wellcome Trust grant (225220/Z/22/Z). E.S., M.S, and M.B. are supported by the Wellcome Trust grant (225220/Z/22/Z). J.R.H. is supported by the Wellcome Trust grants (225220/Z/22/Z and 106130/Z/14/Z) and the MRC grant (MC UU 00029/3).

## Declaration

J.R.H. is a co-founder and director of Nucleome Therapeutics and provides consultancy to the company.

## Footnotes

* For better clarification: *t*_1_ = *H3K4me1, t*_2_ = *H3K4me3, t*_3_ = *H3K27ac*, and *t*_4_ = *CTCF*.

## Notes

### Competing Interest Statement

The authors have declared no competing interest.

